# The inner integument controls embryo sac development and seed shape in *Arabidopsis thaliana*

**DOI:** 10.1101/2024.12.13.628338

**Authors:** Tejasvinee Atul Mody, Kay Schneitz

## Abstract

The angiosperm ovule is characterized by the close association of the two generations, with the haploid female gametophyte or embryo sac being encapsulated by the diploid sporophyte, which usually forms two integuments. How the gametophyte and sporophyte coordinate their development has long been of interest. However, the function of the inner integument in embryo sac development has remained elusive. Here, we addressed this question. We applied a genetic ablation strategy to achieve an early block in inner integument outgrowth. We generated plants expressing *BARNASE* under the control of an early acting endothelium-specific promoter. Corresponding lines carried ovules lacking most of the inner integument. The genetic and cell biological data revealed that in the near absence of an inner integument embryo sac development is blocked at the mono-nuclear embryo sac stage in most pre-fertilization ovules. Approximately 10 percent of the ovules developed a functional embryo sac and underwent fertilization. Subsequent embryo and endosperm development appeared unperturbed and viable seeds were produced albeit of altered shape. Our results show that the inner integument plays an important role in early embryo sac development as well as ovule and seed shape, but is dispensable for embryo and endosperm development. Key words: embryo sac, embryo, endothelium, female gametophyte, integument, seed development

**Highlight:** Genetic ablation of the inner integument demonstrates its role in embryo sac development and its irrelevance for embryogenesis.

## Introduction

The ovule represents the major female reproductive organs of higher plants and is characterized by the close association of the haploid and diploid generations. During its pre-fertilization development, the internal haploid tissue, the female gametophyte or embryo sac, becomes encapsulated by the diploid sporophytic tissue, which contains either one or two integuments. The embryo sac carries the egg cell and the central cell. Seed development starts with fertilization. A double-fertilization event results in the fertilization of the egg and central cells. The fertilized egg cell or zygote develops into the embryo and the fertilized central cell into the endosperm, respectively, while the integuments differentiate into the seed coat.

The complex tissue arrangement in the ovule has led to questions about possible tissue interactions that regulate ovule development. There have been a number of studies on the role of the integuments in these processes, mostly in the model system *Arabidopsis thaliana* (*A. thaliana*) (Gasser *et al*., 1998; Grossniklaus and Schneitz, 1998; Bencivenga *et al*., 2011; Figueiredo and Köhler, 2014; Shirley *et al*., 2019; Doll and Ingram, 2022; Qin *et al*., 2023). However, the mechanisms underlying the perceived crosstalk between these tissues are still largely elusive. A number of sporophytic mutants with a defect in embryo sac development have been found (Bencivenga *et al*., 2011; Qin *et al*., 2023). A prominent example is *inner no outer (ino*). *INO* encodes a transcription factor of the YABBY family and is specifically expressed in the outer layer of the outer integument (Villanueva *et al*., 1999; Balasubramanian and Schneitz, 2000). Loss-of-function alleles of *INO* result in a failure to form the outer integument and the early abortion of embryo sac development indicating an indirect influence of the outer integument on embryo sac development (Baker *et al*., 1997; Schneitz *et al*., 1997; Vijayan *et al*., 2021*a*; Skinner *et al*., 2023). However, more recent results based on quantitative morphometry of a null allele of *INO* revealed defects in the 3D cellular architecture of the internal tissue of the chalaza (Vijayan *et al*., 2021*a*). It raised the possibility that an altered cellular architecture of the chalaza in *ino* mutants could be the result of aberrant cytokinin signaling in the chalaza, which in turn would result in a block in embryo sac development (Vijayan *et al*., 2021*a*). Alternatively, it could impair nutrient delivery (Skinner *et al*., 2023).

The role of the inner integument in the development of the embryo sac, embryo, and endosperm remains unclear as well. The innermost layer of the inner integument develops into the endothelium. It generates an additional cell layer so that the inner integument ultimately consists of three cell layers (Schneitz *et al*., 1995; Debeaujon *et al*., 2003; Vijayan *et al*., 2021*a*). Examining the role of the inner integument in embryo sac and seed development has been difficult. For example, there is a lack of mutants with a specific defect in inner integument development. The homeobox gene *WUSCHEL* (*WUS*) regulates stem cell maintenance in the shoot apical meristem and floral meristem (Mayer *et al*., 1998; Brand *et al*., 2000; Schoof *et al*., 2000).

Interestingly, *wus-7*, a weak allele frequently causes a reduced inner integument and a defective embryo sac (Lora *et al*., 2019). However, *WUS* is expressed in the nucellus, the sporophytic tissue that contains the developing embryo sac, and the early inner integument (Gross-Hardt *et al*., 2002; Sieber *et al*., 2004; Vijayan *et al*., 2021*a*). Moreover, *WUS* is required for early megasporogenesis (Lieber *et al*., 2011) and for the formation of the chalaza and both integuments (Gross-Hardt *et al*., 2002; Sieber *et al*., 2004). Thus, despite the reduced inner integument in *wus-7* it remained unclear whether the inner integument influences the development of the female gametophyte.

During seed development, the endothelium produces proanthocyanidins (PAs), condensed tannins, which are required for seed dormancy and germination (Devic *et al*., 1999; Debeaujon *et al*., 2000, 2001). *ARABIDOPSIS B SISTER*/*TRANSPARENT TESTA16* (*ABS*/*TT6*) encodes a MADS domain transcription factor that controls endothelium differentiation and PA biosynthesis (Nesi *et al*., 2002). Despite those alterations, *abs*/*tt16* mutants produced viable seeds. *SEEDSTICK* (*STK*) encodes another MADS domain transcription factor (Pinyopich *et al*., 2003) that can form protein complexes with ABS/TT16 (Kaufmann *et al*., 2005)*. ABS/TT16* and *STK* are both specifically expressed in the endothelium of ovules and *abs stk* double mutants exhibit failed differentiation in the innermost integument layer, smaller embryo sacs, failure of polar nuclei fusion in the central cell, ectopic starch accumulation in the embryo sac, fertilization defects, and delayed embryo development (Mizzotti *et al*., 2012). Genetic ablation of the endothelium during seed development had no appreciable effect on embryo and endosperm development but resulted in reduced seed dormancy (Debeaujon *et al*., 2003). This result indicates that the endothelial cell layer of the inner integument is not essential for the development of both tissues.

Here, we address the role of the inner integument in ovule and seed development by specifically blocking early inner integument development using a tissue-specific cell ablation strategy. Our data indicate that the inner integument plays an important role in embryo sac development and seed shape but is not required for the development of viable seeds and functional seedlings.

## Materials and methods

### Plant work and growth conditions

*Arabidopsis thaliana* (L.) Heynh. var. Columbia (Col-0) was used as the wild-type strain. Col-0 and *pTT1::lt-BAR* plants were grown as described earlier (<u>Fulton et al., 2009</u>). The FGR8.0 line (Völz *et al*., 2013) and the pPHE1::PHE1-GFP line (Weinhofer *et al*., 2010) were described earlier.

### DNA work and plant transformation

For DNA work, standard molecular biology techniques were used. PCR fragments used for cloning were obtained using Phusion DNA polymerase (New England Biolabs, Frankfurt, Germany). All PCR-based constructs were sequenced. Constructs were generated with the help of the GreenGate system (Lampropoulos *et al*., 2013).

The *pTT1::ltBAR:tTT1* construct was generated as follows. The *3kbProTT1:gTT1* plasmid (Coen *et al*., 2019) was used as a template to amplify the 3.1 kb *pTT1*. The 3.1 kb *pTT1* contained an internal BsaI site which would hinder cloning using the greengate system. Therefore, in a first step the internal BsaI site was removed and then the *pTT1* fragment was ligated to the pGGA greengate entry vector. The following primers were used for *pTT1* amplification with BsaI overhangs and for removing the internal BsaI restriction site: pTT1 (3.1 Kb) Fw BsaI (5’-AACAGGTCTCAACCTAACCATTTGCTTGTGTCAACAACAT-3’), pTT1 (3.1 Kb) Rw BsaI (5’-AACAGGTCTCTTGTTTGAATGTGGTGAATAGTTGTTGGAG-3’), pTT1 (3.1 Kb) F0 BsaI (5’-AACAGGTCTCTGTCCCGGCATAATATTTGACTAGGC-3’), and pTT1 (3.1 Kb) R0 BsaI (5’-AACAGGTCTCGGGACTCCACAATTCAATATTCATA-3’). The 5.8 kb pGGA-pTT1 included 2.6 kb plasmid backbone and 3.1 kb *pTT1* insert. The 900 bp *tTT1* was amplified using the primers: tTT1 Rw-bsa1 (5’-AACAGGTCTCTTAGTAAAAATTCTAATGATTTGTTTTTTTCC-3’) and tTT1 fw-bsa1 (5’-AACAGGTCTCACTGCATTTGGGCATCTTTTTCTTT-3’). The 3.5 kb pGGE-tTT1 plasmid was also generated using GreenGate cloning. The pENTR::lt-BAR plasmid was a gift from Nico Dissmeyer. The lt-BAR construct (Faden et al., 2019) also had an internal BsaI site. The following primers were used for lt-BAR amplification with BsaI overhangs and for removing the internal BsaI restriction site: K2barn_R0 (5’-AACAGGTCTCAGTCCCCGTTCTTGCCAATC-3’), K2barn_F0 (5’-AACAGGTCTCAGGACCTACCCTGGCCTCCGC-3’), K2barn_BsaI_RW (5’-AACAGGTCTCTCTGATCTGATTTTTGTAAAGGTCTG-3’), and K2barn_BsaI_FW (5’-AACAGGTCTCAGGCTCAACATTTGTACAAAAAAGCAGGCT-3’). The about 4 kb pGGC-lt-BAR plasmid was assembled again using GreenGate cloning. The pGGZ-*pTT1::It-BAR:tTT1* was assembled using the standard GreenGate reaction involving ligation of the GreenGate cloning entry vector parts: pGGA-pTT1, pGGB003, pGGC-lt-BAR, pGGD002, pGGE-tTT1, pGGF007 (KanR), and the pGGZ001 destination vector. All PCR-based constructs were sequenced. Arabidopsis Col-0 wild-type plants were transformed with the *pTT1::lt-BAR* construct using Agrobacterium strain GV3101/pMP90 (Koncz and Schell, 1986) and the floral dip method (Clough and Bent, 1998). Transgenic T1 plants were selected on Kanamycin (50 μg/mL) plates and transferred to soil for further inspection.

The *pTT1::nls:GFP* line and construct are an abbreviation of *pTT1-lox-nlsGFP-rbcSt-lox-PME5-tTT1*. Since this line lacks a Cre construct, only *pTT1::nls:GFP* is expressed, hence the abbreviation. The *pTT1::nls:GFP* construct was generated as follows. The pGGB-*lox-nlsGFP-rbcSt-lox* plasmid was assembled using GreenGate cloning. The lox-nlsGFP-rbcS-lox fragment was amplified from a pGGM-pINO-nlsGFP-rbcS plasmid using primers: Fwd primer (GGC-lox-GFP fwd: 5’-AACAGGTCTCAAACAATAACTTCGTATAGCATACATTATACGAAGTTATATGGAAGAGCAAGCAAGGAAAGC-3’) and Rev primer (GGC-lox-rbcs rev: 5’-AACAGGTCTCTAGCCATAACTTCGTATAGCATACATTATACGAAGTTATCGATTGATGCATGTTGTCAATC-3’). The pGGC-PME5 plasmid was assembled using GreenGate cloning after amplifying the coding sequence of *PME5* using Col-0 cDNA as template and using the primers: pGGC-PME5 fwd (5’-AACAGGTCTCAGGCTCAACAATGGCGCAACTTACTAATTC-3’) and pGGC-PME5rev (5’-AACAGGTCTCTCTGATTAAGCATCTCGAGGAGCG-3’). The pGGZ-*pTT1-lox-nlsGFP-rbcSt-lox-PME5-tTT1* was assembled using the standard GreenGate reaction involving ligation of the GreenGate cloning entry vector parts: pGGA-pTT1, pGGB-*lox-nlsGFP-rbcSt-lox*, pGGC-PME5, pGGD002, pGGE-tTT1, pGGF005 (HygR), and the pGGZ001 destination vector. All PCR-based constructs were sequenced. Arabidopsis Col-0 wild-type plants were transformed with the pGGZ-*pTT1-lox-nlsGFP-rbcSt-lox-PME5-tTT1* construct using Agrobacterium strain GV3101/pMP90 (Koncz and Schell, 1986) and the floral dip method (Clough and Bent, 1998). Transgenic T1 plants were selected on Hygromycin (20 μg/mL) plates and transferred to soil for further inspection.

### Clearing and staining of tissue samples

Treatment of ovules and seeds was done as described in (Tofanelli *et al*., 2019) and (Vijayan *et al*., 2021*b*) with some optimizations. Tissue was fixed in 4% paraformaldehyde in PBS for 1-1.5 h followed by one wash in PBS before transfer into the ClearSee solution (xylitol (10%, w/v), sodium deoxycholate (15%, w/v), urea (25%, w/v), in H_2_O) (Kurihara *et al*., 2015). Clearing was done at least overnight or for up to two days. Cell wall staining with SR2200 (Renaissance Chemicals, Selby, UK) was performed as described in (Musielak *et al*., 2015). Cleared tissue was washed in a PBS solution containing 0.1% SR2200 and then put into a PBS solution containing 0.1% SR2200 and a 1/1000 dilution of the nuclear stain TO-PRO-3 iodide (Thermo Fisher Scientific) for 30 minutes. Tissue was washed in PBS for one minute, transferred again to ClearSee for 20 minutes before mounting in Vectashield antifade agent (Vector Laboratories, Burlingame, CA, USA).

### Microscopy and image acquisition

Confocal laser scanning microscopy of ovules, seeds, and embryos stained with SR2200 and TO-PRO-3 iodide was performed on an upright Leica TCS SP8 X WLL2 HyVolution 2 (Leica Microsystems) equipped with GaAsP (HyD) detectors and a 63x glycerol objective (HC PL anterior-posteriorO CS2 63x/1.30 GLYC, CORR CS2). Scan speed was at 400 Hz, the pinhole was set to 1 Airy units, line average between 2 and 4, and the digital zoom between 0.75 and 2. For z-stacks, 8, 12, or 16 bit images were captured at a slice interval of 0.33 μm with voxel size of 0.126 μm x 0.126 μm x 0.33 μm. Laser power or gain was adjusted for z compensation to obtain an optimal z-stack. Images were adjusted for color and contrast using Adobe Photoshop 2021 (Adobe, San Jose, USA) or MorphographX software (Strauss *et al*., 2022). Image acquisition parameters were the following: SR2200; 405 diode laser 0.10%, HyD 416–476 nm, detector gain 10. TO-PRO-3; 642 nm White Laser 2%, HyD 661–800 nm, detector gain 400. Image acquisition parameters for the pPHE1::PHE1-GFP reporter line were the following: SR2200; 405 diode laser, HyD 416–476 nm. GFP; 488 nm Argon laser, HyD 493–626 nm. TO-PRO-3; 642 nm White Laser, HyD 660– 800 nm. Image acquisition parameters for the FGR8.0 reporter line were the following: SR2200; 405 diode laser, HyD 416–476 nm. GFP and YFP; 488 and 514 nm Argon laser, PMT 524–553 nm. DsRed; 558 nm White Laser, HyD 575–731 nm. The GFP and YFP emissions were detected together because it was not possible to exclude the 3XGFP emission from the YFP channel. In each case, sequential scanning was performed to avoid crosstalk between the spectra.

### Datasets, 3D cell segmentation, and 3D cell meshes

The dataset encompassing the segmented wild-type 3D digital ovules of *Arabidopsis thaliana* was described earlier (Vijayan *et al*., 2021*b*). The z-stacks of *pTT1::BAR* ovules were 3D cell segmented using the PlantSeg pipeline (Wolny *et al*., 2020). In all instances cell 3D meshes were generated with MorphoGraphX using segmented image stacks using the process “Mesh/Creation/Marching Cube 3D” with a cube size of 1. Manual cell type labeling was performed with MorphoGraphX. The full *pTT1::BAR* ovule dataset, including raw cell boundaries, cell boundaries, PlantSeg predictions, nuclear images, segmented cells, and the annotated 3D cell meshes, as well as the associated attribute files in csv format, can be found in the Biostudies data repository at EMBL-EBI (Sarkans *et al*., 2018) (https://www.ebi.ac.uk/biostudies) (accession S-BIAD1499). The Col-0 wild-type embryo 3D meshes were obtained from (Yoshida *et al*., 2014). The 3D mesh files can be opened in MorphoGraphX. The equivalent Arabidopsis wild-type ovule dataset on Biostudies has the accession S-BSST475.

### Exporting attributes from MorphoGraphX for further quantitative analysis

All quantitative cellular features were exported as attributes from MGX. The attributes included cell IDs (segmentation label of individual cells), cell type IDs (tissue annotation), and cell volume. The attributes from individual ovules were exported as csv files and merged to create long-format Excel-sheets listing all the scored attributes of all the cells from the analyzed ovules. *pTT1::BAR* ovule cell IDs with volume less than 20 μm^3^ have been excluded from cellular analyses since these correspond to artifacts and empty spaces that are segmented as cells. The files are included in the downloadable datasets.

### Software

The MorphographX software was used for the generation of cell 3D and surface meshes, cell type labeling, and the analysis of 3D cellular features (Barbier de Reuille *et al*., 2015; Strauss *et al*., 2022). It can be downloaded from its website (https://morphographx.org). The PlantSeg pipeline (Wolny *et al*., 2020) was used for 3D cell boundary prediction and segmentation. The software can be obtained from its Github repository (https://github.com/kreshuklab/plant-seg).

### Statistical analysis

Statistical analysis was performed using a combination of R (R Development Core Team, 2024; https://www.R-project.org/), RStudio (R Studio Team, 2024; https://posit.co/), Tidyverse (Wickham *et al*., 2019), and PRISM10 software (GraphPad Software, San Diego, USA).

#### Box and whiskers plots

Boxplots show the median value of the distribution as a central line and mean value of the distribution as a plus sign within the box. The limits of the box represent the quartiles of the distribution. Whisker ends mark the minimum and maximum of all the data.

## Results

### Ovule development in *Arabidopsis thaliana*

The *Arabidopsis thaliana* ovules emerge as finger-like protrusions from the placenta and after tissue differentiation, develop a curved shape visible in mature ovules and also in seeds (Fig. 1A,B). During stage 2 of ovule development, megasporogenesis in the distal nucellus originates the functional megaspore that will develop into the haploid embryo sac during stage 3. The inner integument (ii) and outer integument (oi) emerge laterally from the central chalaza. Each integument is composed of two cell layers, an inner (ii1 and oi1) and an outer layer (ii2 and oi2). The ii1 layer is also known as endothelium. Each of these layers is of single cell thickness (Fig. 1B). In stage 3 ovules, the endothelium (ii1) originates an additional cell layer (ii1’) by periclinal asymmetric cell divisions. During stage 3, the two integuments grow asymmetrically over the nucellus, and eventually the ovule exhibits a pronounced curvature (Schneitz *et al*., 1995; Vijayan *et al*., 2021*b*).

**Fig. 1.**
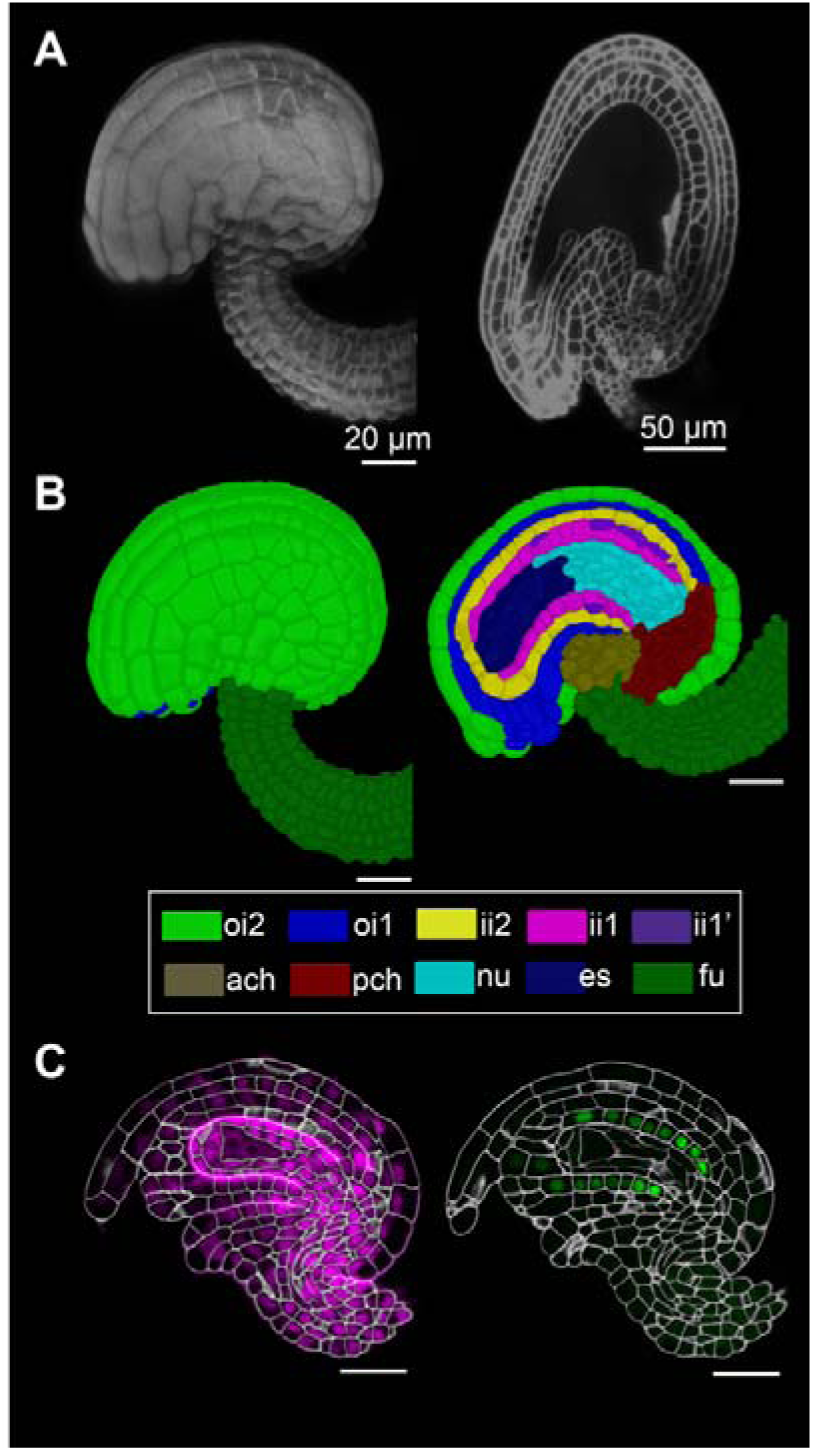
Tissue organization in the wild-type ovule and *pTT1::nls:GFP* expression in the inner layer of inner integument. A. Left: 3D rendering of confocal z-stack of SR2200-stained cell walls of *Arabidopsis thaliana* Col-0 stage 3-V ovule; right: mid-sagittal section of 3D rendering of confocal z-stack of SR2200-stained cell walls of *Arabidopsis thaliana* Col-0 seed harboring a 2-celled embryo. B. 3D view (left) and mid-sagittal section (right) of a stage 3-V ovule with cell-type labels and showing the cell type organization in wild-type ovules. C. Mid-sagittal section of a stage 3-III *Arabidopsis thaliana* ovule expressing *pTT1::nls:GFP*. The cell outlines shown are cell boundary predictions obtained from SR2200-stained cell walls. Left: TO-PRO-3-stained nuclei (magenta) along with cell outlines and right: *pTT1::nls:GFP* signal in the inner layer of inner integument. Abbreviations: ach, anterior chalaza; es, embryo sac; fu, funiculus; ii1, inner layer of inner integument; ii1’, parenchymatic layer of inner integument; ii2, outer layer of inner integument; nu, nucellus; oi1, inner layer of outer integument; oi2, outer layer of outer integument; pch, posterior chalaza. Scale bars: A, left: 20 μm and right: 50 μm; B and C, 20 μm.

### Generation of a mutant with reduced inner integument

To obtain mutant lines carrying ovules with nearly eliminated inner integuments we generated transgenic lines harboring the cytotoxic bacterial RNase barnase gene under the control of 3kb promoter of the *TRANSPARENT TESTA 1* (*TT1*). *TT1* encodes a C2H2 zinc finger transcription factor and controls endothelium development and ii1’ polar cell patterning (Sagasser *et al*., 2002; Coen *et al*., 2019). A 3kb genomic fragment of the *TT1* promoter was reported to drive *TT1* expression specifically in the endothelium (ii1) in Arabidopsis ovules (Coen *et al*., 2019). To confirm the tissue specificity of the 3kb *TT1* promoter fragment we generated transgenic reporter lines harboring a construct where this promoter fragment drives expression of GFP fused to a nuclear localization signal (*pTT1::nlsGFP*) (see Materials and Methods). To investigate the expression pattern of this reporter we performed 3D confocal laser scanning microscopy of fixed and cleared ovules from several transgenic lines that were also stained with the cell wall stain SR2200 and nuclear stain TO-PRO-3. We scored ovules from the primordia stage (stage 1) up to stage before fertilization (stage 3-VI). No expression was detected in ovule primordia. Starting from stage 2-III to 2-V ovules, expression becomes visible only in a few, but not all cells of the developing endothelium. During stage 3 we detected prominent and specific *pTT1* expression in the endothelium and the ii1’ layer (Fig. 1C). In summary, our data suggest endothelium-specific expression of the *pTT1::nlsGFP* reporter confirming previous results (Coen *et al*., 2019).

Next, we performed genetic cell ablation experiments. Expression of toxic proteins, such as bacterial RNase barnase (BAR) is a powerful approach to eliminate entire cell populations or specific tissues in a biological context (Buckle and Fersht, 1994; Leuchtenberger *et al*., 2001; Lännenpää *et al*., 2005). The BAR toxin has been shown previously to cause cell growth arrest rather than cell death in trichome cells (Faden *et al*., 2019). The low temperature (lt) degron when combined with BAR, can enable cell ablation in a temperature-dependent (conditional) manner and prevent the reliance on exogenous compounds such as hormones or other small molecules for conditional, non-constitutive, cell ablation (Faden *et al*., 2016; Dissmeyer, 2017). We reasoned that expressing *lt-BAR* under the control of the 3kb *TT1* promoter fragment should result in the conditional ablation of the inner integument. Therefore, we generated more than 20 independent transgenic *pTT1::lt-BAR* lines in the *Arabidopsis thaliana* Col-0 background. We analyzed ovule phenotype of 5 transgenic lines in more detail. Under our growth conditions we observed essentially identical defects in inner integument development in those 5 lines irrespective of whether the plants were grown under restrictive or permissive temperature conditions. These findings suggest that the lt-degron seemed non-functional or leaky in our experiments. Hence, from now on we refer to this *pTT1::lt-BAR* construct as *pTT1::BAR*.

To investigate the impact of BAR activity on the inner integument of *pTT1::BAR* ovules, we first generated 3D digital ovules of two independent *pTT1::BAR* lines (lines 9 and 10). To this end, we performed 3D confocal laser scanning microscopy (CLSM) of fixed, cleared, and stained ovules to obtain z-stacks of ovules. This was followed by cell boundary prediction and 3D cell segmentation using the PlantSeg pipeline. The segmented stacks pertaining to ovules of line 9 were then loaded into the MorphographX (MGX) software for mesh creation, cell-type labeling, and quantitative analyses (see Materials and Methods). We generated over 50 cell-type labeled stage 3 3D digital *pTT1::BAR* ovules. The *pTT1::BAR* ovule is visually characterized by a reduction in inner integument development (Fig. 2A) in comparison to the Col-0 wildtype ovules (Fig. 1B). It is also important to note that there is variability in the morphology and reduction in inner integument development of *pTT1::BAR* ovules (Fig. S1). The inner integument reduction encompasses the endothelial (ii1), ii1’ and ii2 layers. Therefore, it seems that an impact on the ii1 layer also affects the ii2 layer of the inner integument.

**Fig. 2.**
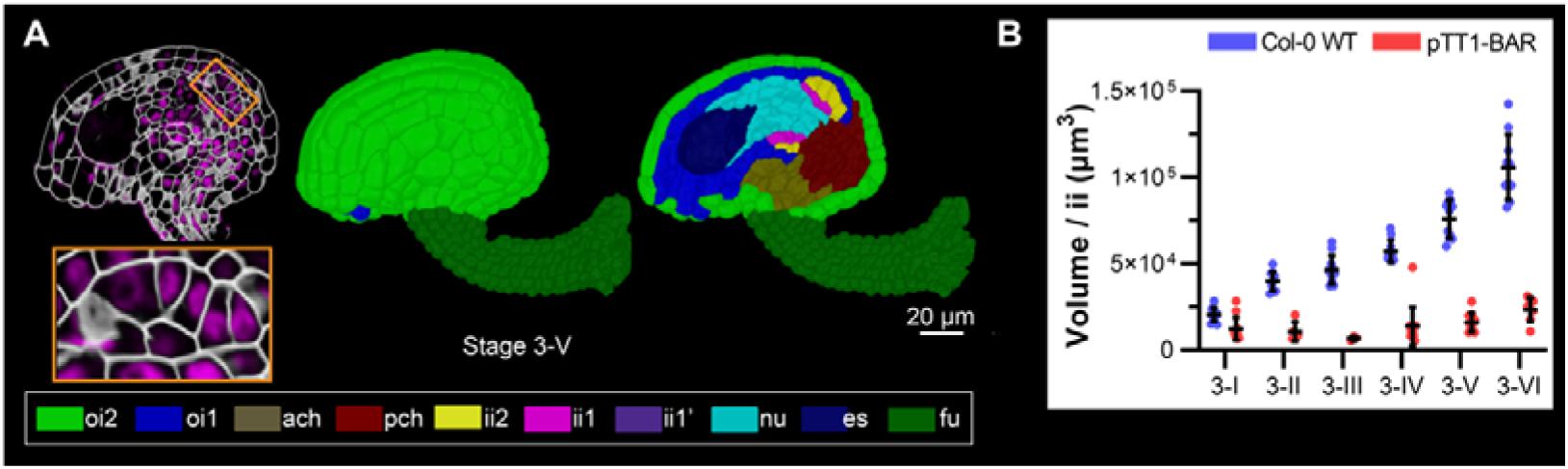
*Arabidopsis thaliana pTT1::BAR* ovules demonstrate a reduction in inner integument growth. A. Stage 3-V *Arabidopsis thaliana pTT1::BAR* ovule. A. left: mid-sagittal section of *pTT1::BAR* ovule showing cell-outlines along with TO-PRO-3-stained nuclei (magenta). The orange box is used to demonstrate a zoomed-in inner integument below to visualize the intact nuclear morphology of inner integument cells. A. middle: 3D view and right: mid-sagittal section (right) of a stage 3-V *pTT1::BAR* ovule with cell-type labels showing the tissue organization. B. Plot depicting the volume of inner integument of WT and *pTT1::BAR* ovules from stages 3-I to 3-VI of ovule development. Data points indicate individual ovules. Mean ± SD is shown. Abbreviations: ach, anterior chalaza; es, embryo sac; fu, funiculus; ii1, inner layer of inner integument; ii1’, parenchymatic layer of inner integument; ii2, outer layer of inner integument; nu, nucellus; oi1, inner layer of outer integument; oi2, outer layer of outer integument; pch, posterior chalaza. Scale bar: 20 μm.

To quantify the reduction in inner integument growth in terms of volume, we quantified and compared the inner integument volumes and cell numbers of stage 3 *pTT1::BAR* ovules of line 9 with Col-0 wild-type ovules (data from (Vijayan *et al*., 2021*b*) using MGX (see Materials and Methods for details). We observed that the stage 3 *pTT1:BAR* ovules showed an arrest in the increase of the inner integument volume compared to the Col-0 wild-type ovules. The reduction in volume of *pTT1::BAR* ovules was more pronounced in late stage 3 ovules than in early stage 3 ovules (Fig. 2B, Table 1). The volumes of stage 3-I and 3-V *pTT1::BAR* ii are 1.22 × 10^4^ ± 0.66 × 10^4^ µm^3^ (mean ± SD)) and 1.59 × 10^4^ ± 0.54 × 10^4^ µm^3^, respectively; whereas, those of the ii of wild type were 2.05 × 10^4^ ± 0.41 × 10^4^ µm^3^ and 7.59 × 10^4^ ± 1.11 × 10^4^ µm^3^ respectively. This translates to a reduction in ii volume of *pTT1::BAR* ovules of approximately 40% at stage 3-I and 80% at stage 3-V compared to wild-type ovules of the same stage. Table 1 shows the total ii volumes for *pTT1::BAR* and wild-type ovules along with the percentages of ii volume reduction in *pTT1::BAR* ovules.

**Table 1.**
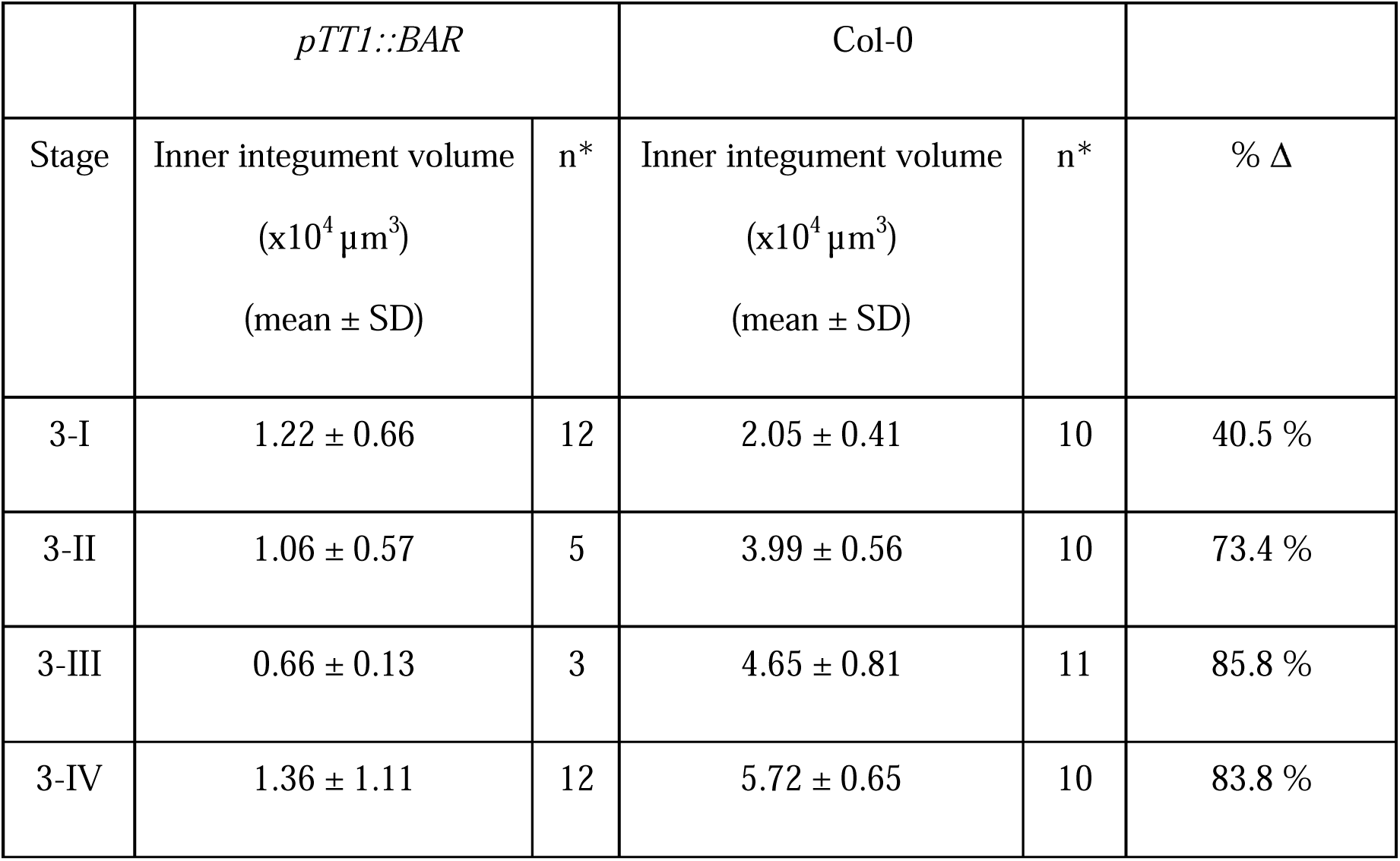
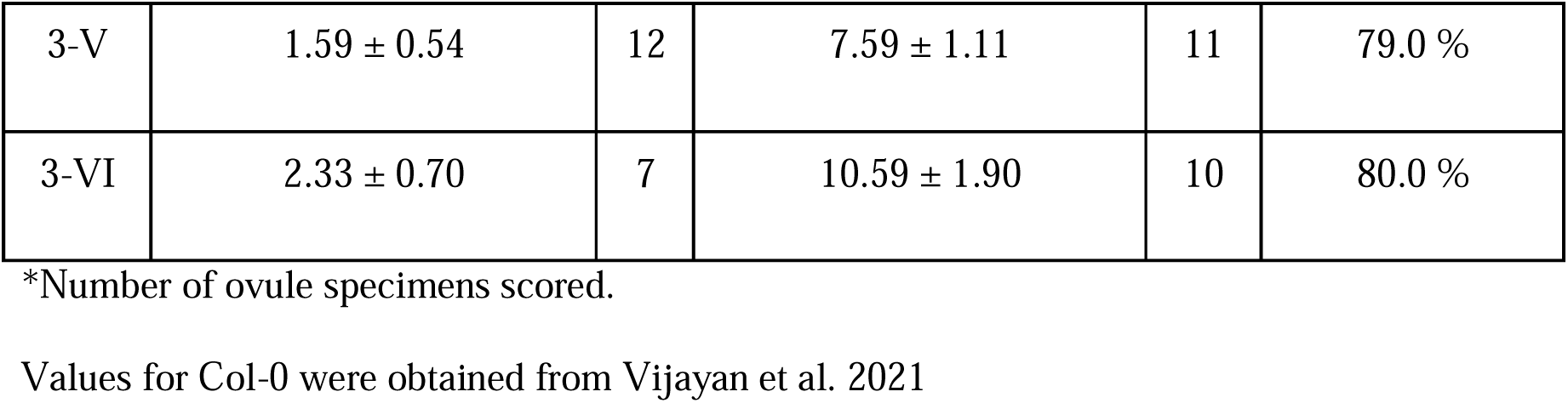
Stagewise volume of inner integument of *pTT1::BAR* and wild-type ovules.

The total cell number of stage 3-I and 3-V *pTT1::BAR* ii were 63 ± 28 (mean ± SD) and 67 ± 18, respectively; whereas, that of the ii of wild type is 132 ± 18 and 343 ± 48, respectively. This implies a reduction in ii cell number of *pTT1::BAR* ovules by about 52% at stage 3-I and 80% at stage 3-V. The nuclear morphology of ii cells of *pTT1::BAR* ovules seemed to remain intact, as shown by TO-PRO-3 staining of nuclei (Fig. 2A). Together, our observations of reduced ii tissue volume, ii cell number, and intact ii nuclear morphology in *pTT1::BAR* ovules suggest that *BAR* activity in the ii layer causes cell proliferation arrest and tissue growth arrest but not cell death in the inner integument.

### Reduction of inner integument growth affects the development of other ovule tissues

To identify and test for any effects of a reduced inner integument on the growth of other sporophytic tissues of the ovule, we compared individual tissue volumes (Table 2), cell numbers (Table 3), and cell volumes (Table 4) of *pTT1::BAR* and Col-0 wild-type ovules at stages 3-I and 3-V (Fig. 3). We found that the size of the chalaza, embryo sac, and funiculus remained largely unaffected. There were less than 10% and statistically non-significant differences in tissue volume, cell volume, and cell number between mutant and wild-type ovules. However, in stage 3-V *pTT1::BAR* ovules, the size of the outer integument was reduced by approximately 15% compared to wild-type ovules. The stage 3-V *pTT1::BAR* outer integument was also characterized by an 18% reduction in cell number and a 3% increase in average cell volume compared to wild type. In contrast, the stage 3-V *pTT1::BAR* ovule nucellus showed a 34% increase in volume and 23% more cells, which were approximately 8% larger than wild type. Overall, the reduction of the inner integument primarily affects cell proliferation and, to a lesser extent, cell growth, of the adjacent outer integument and nucellus.

**Table 2.** Tissue volumes of stages 3-I and 3-V ovules of *pTT1::BAR* and wild type.

|  | *pTT1::BAR* |  | Col-0 |  |
| --- | --- | --- | --- | --- |
| Tissue volumes<br>(x10<sup>4</sup> μm<sup>3</sup>)<br>(mean ± SD) | 3-I<br>(n*=12) | 3-V<br>(n*=12) | 3-I<br>(n*=10) | 3-V<br>(n*=11) |
| Inner integument | 1.22 ± 0.66 | 1.59 ± 0.54 | 2.05 ± 0.41 | 7.59 ± 1.11 |
| Outer integument | 12.05 ± 2.72 | 15.74 ± 1.66 | 8.77 ± 1.72 | 18.57 ± 1.32 |
| chalaza | 3.11 ± 0.96 | 4.24 ± 0.78 | 1.92 ± 0.44 | 3.91 ± 0.69 |
| nucellus | 1.17 ± 0.52 | 1.58 ± 0.29 | 0.70 ± 0.12 | 1.18 ± 0.27 |
| Embryo sac | 0.08 ± 0.05 | 1.11 ± 0.32 | 0.17 ± 0.09 | 1.05 ± 0.26 |
| funiculus | 4.41 ± 1.39 | 6.98 ± 1.12 | 4.46 ± 0.55 | 7.10 ± 0.83 |

\*Number of ovule specimens scored.
Values for Col-0 were obtained from Vijayan et al. 2021

**Table 3.**
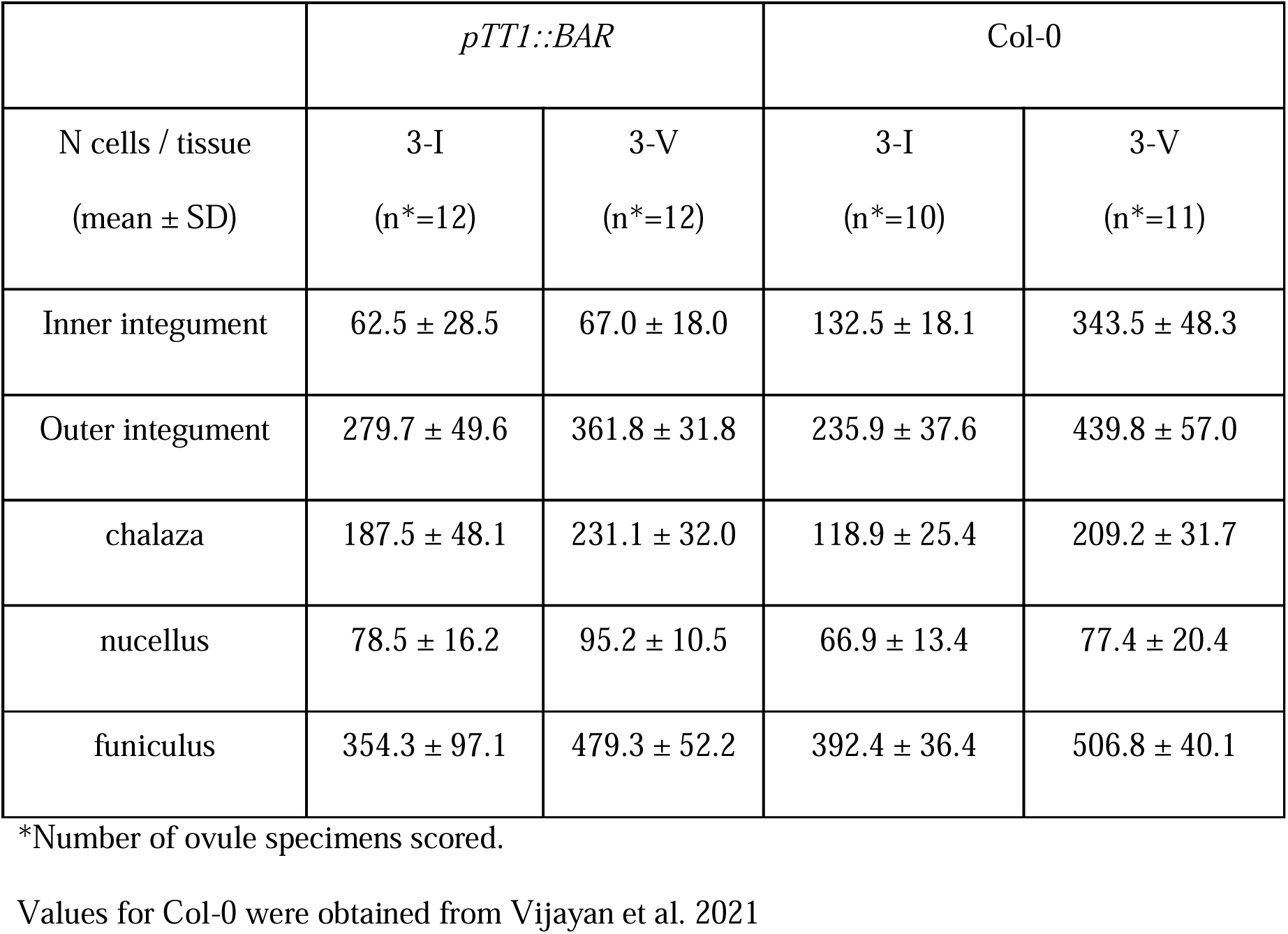
Tissue cell numbers of stages 3-I and 3-V ovules of *pTT1::BAR* and wild type.

**Table 4.** Cell volumes of stages 3-I and 3-V ovules of *pTT1::BAR* and wild type.

|  | *pTT1::BAR* |  | Col-0 |  |
| --- | --- | --- | --- | --- |
| Cell volume (µm<sup>3</sup>)<br>(mean ± SD) | 3-I<br>(n*=12) | 3-V<br>(n*=12) | 3-I<br>(n*=10) | 3-V<br>(n*=11) |
| Inner integument | 194.9 ± 123.3 | 236.7 ± 155.8 | 154.3 ± 74.6 | 220.9 ± 120.2 |
| Outer integument | 430.7 ± 310.9 | 435.0 ± 294.9 | 371.6 ± 244.1 | 422.2 ± 327.6 |
| chalaza | 165.5 ± 124.4 | 183 ± 142.5 | 161.6 ± 105.0 | 187.0 ± 166.5 |
| nucellus | 148.9 ± 98.4 | 165.2 ± 97.5 | 104.3 ± 88.8 | 152.2 ± 204.5 |
| funiculus | 123.4 ± 51.0 | 144.5 ± 69.5 | 113.8 ± 46.4 | 140.1 ± 60.7 |

\*Number of ovule specimens scored.
Values for Col-0 were obtained from Vijayan et al. 2021

**Fig. 3.**
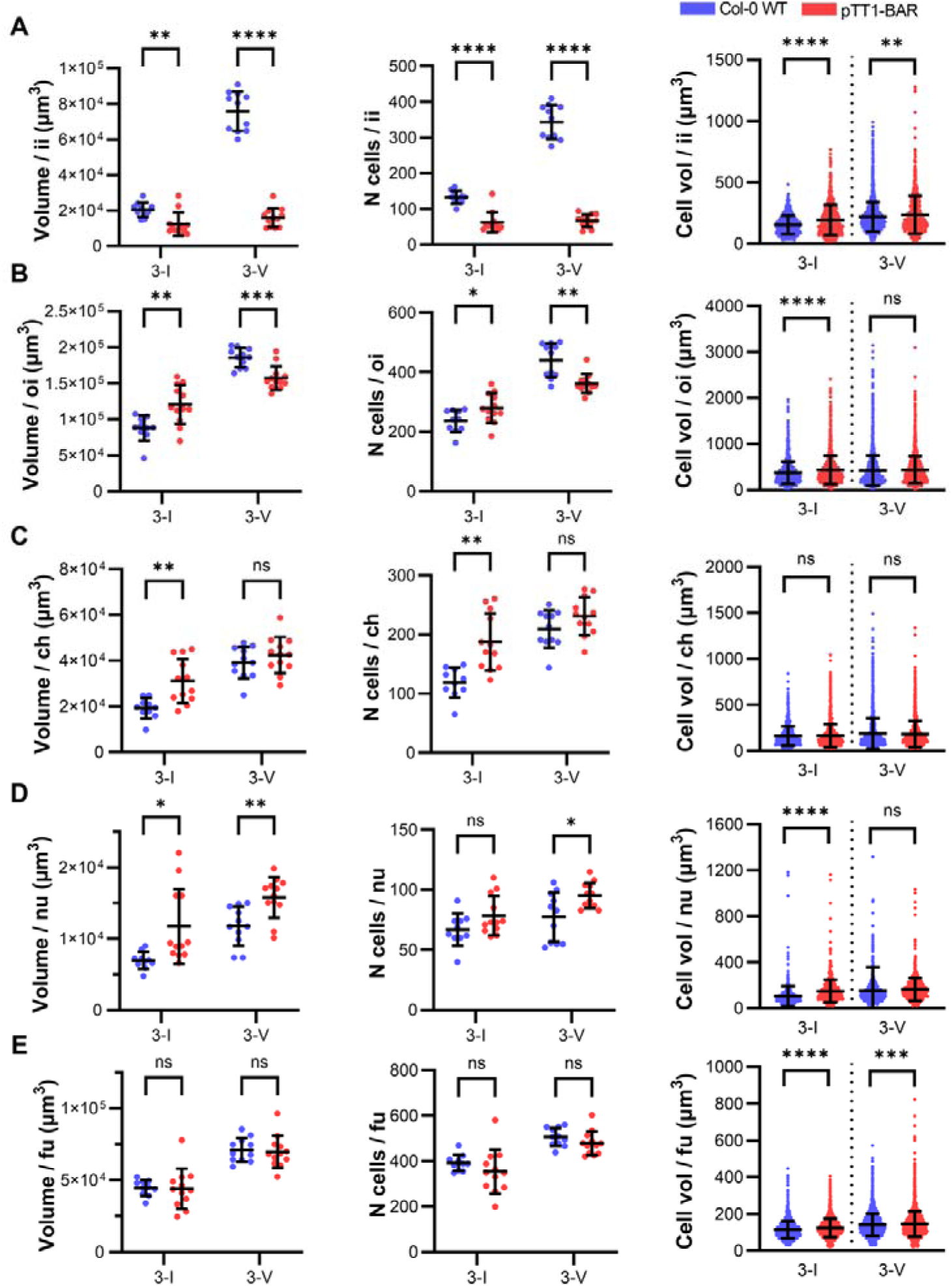
Effect of inner integument reduction on the growth of ovule tissues. Plots depicting a comparison of tissue-specific volumes (left panels), cell numbers (middle panel), and cell volumes between WT and *pTT1::BAR* stage 3-I and 3-V ovules. A. Inner integument, B. Outer integument, C. Chalaza, D. Nucellus, and E. Funiculus. Data points indicate individual ovules. Mean ± SD is shown. Stages are indicated on the x-axes.

### The inner integument demonstrates a scaffolding function in the ovule by influencing ovule shape

In addition to the effects of reduced integument growth on the growth and cellular properties of other tissues, we investigated the structural importance of the integument for the shape of the ovule. Specifically, we asked how a reduced inner integument affects the form of other tissues. To this end, we examined the overall morphology and curvature of the entire ovule, as well as that of the outer integument and the nucellus of *pTT1::BAR* and wild-type ovules. We observed a marked difference in form between stage 3-V ovules of mutant and wild-type (Fig. 1B, 2A, and S1). The outer and inner layers of the outer integument of stage 3-V *pTT1::BAR* ovules exhibited a sharper rather than a more gradual bend in the distal region. This was more pronounced in the inner layer of the outer integument (Figure 4A-C). In addition, the *pTT1::BAR* outer integument, when viewed from the proximal end, appeared to have a tapered phenotype at the distal end compared to the wild type, where no such taper was evident (Fig. 4D).

**Fig. 4.**
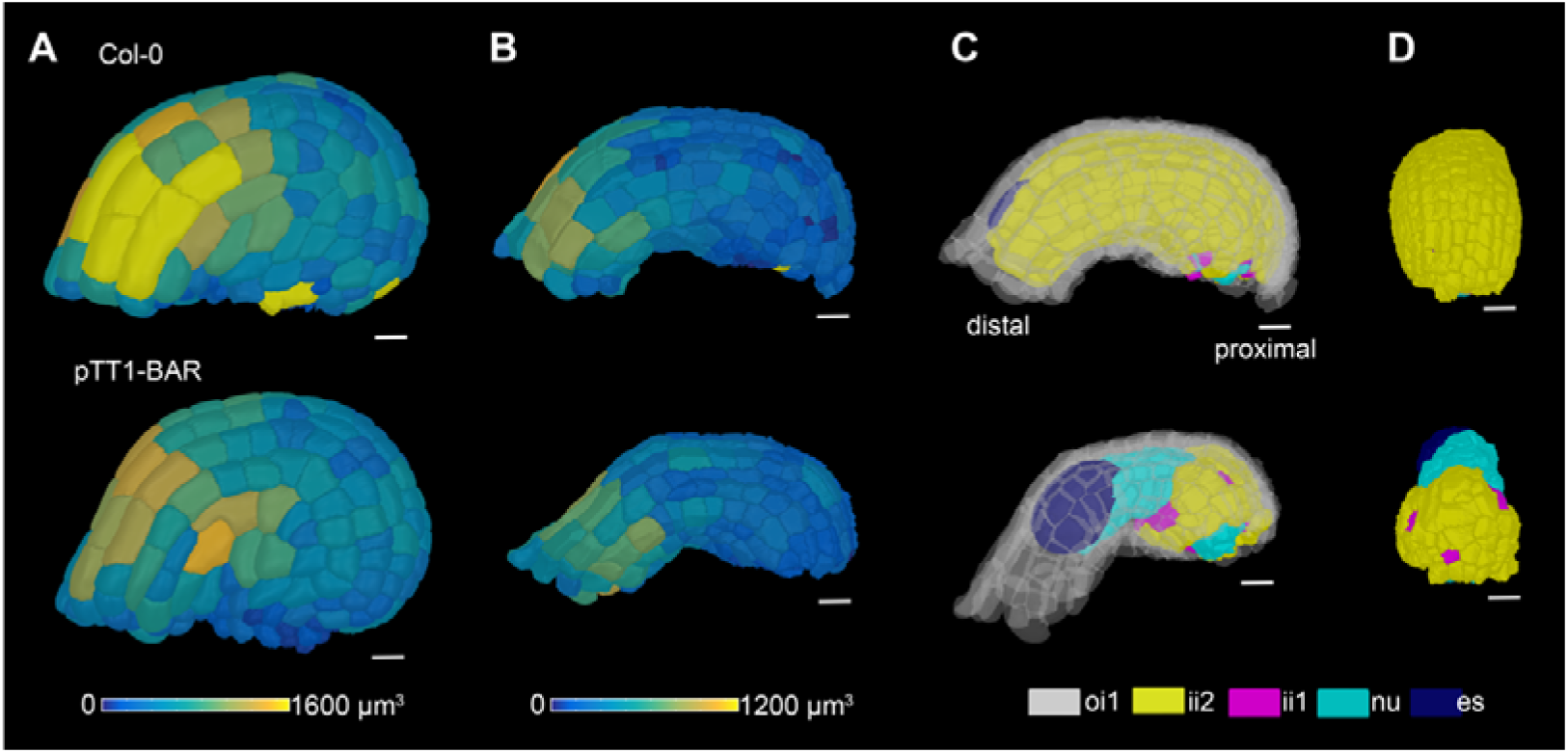
The inner integument demonstrates a scaffolding function in the ovule. Comparison of integument morphology of Col-0 WT (top panel) and *pTT1::BAR* (bottom panel) ovules. A. 3D cell mesh of the outer layer of the outer integument (oi2) showing heat maps of the cell volume in the range from 0 to 1600 μm^3^. B. 3D cell mesh of the inner layer of the outer integument (oi1) showing heat maps of the cell volume in the range from 0 to 1200 μm^3^. C. 3D cell meshes showing oi1 along with the tissues underneath including the two layers of the inner integument, nucellus, and the embryo sac. The Col-0 oi1 is gradually curved (top), whereas the *pTT1::BAR* oi1 is more sharply bent at the distal end. D. The Col-0 inner integument growth is complete from proximal to distal end. In *pTT1::BAR* ovules the reduced inner integument is present only at the proximal end and lacking distal growth. Scale bars: 10 μm.

Despite their size differences, the nucelli of *pTT1::BAR* and wild-type ovules were curved (Figure 5A), suggesting that the outer integument is sufficient to impose curvature on the nucellus when the inner integument is absent or reduced. Taken together, these results suggest that the inner integument contributes to ovule curvature mainly by influencing the development and form of the outer integument. Thus, we hypothesize that the inner integument has a scaffolding function in the ovule, influencing the proper development of the outer integument and the subsequent overall shape and curvature of the ovule.

**Fig. 5.**
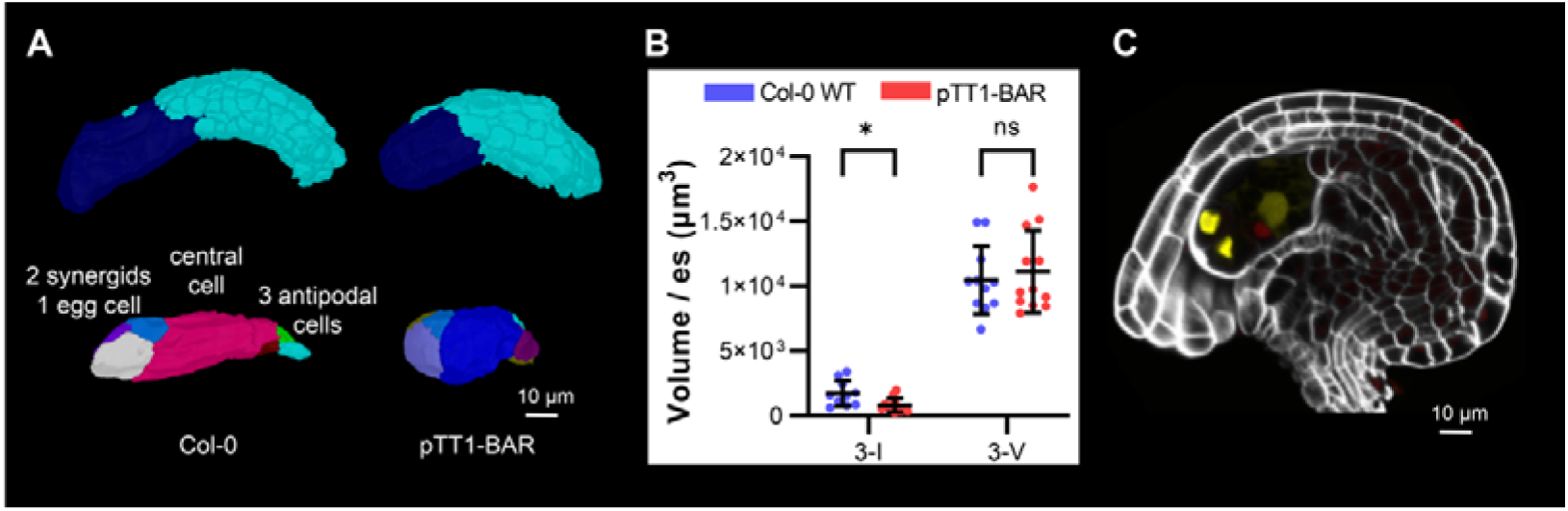
Embryo sac morphology of Col-0 WT and *pTT1::BAR* stage 3-V ovules. A. Top: Nucellus (cyan) and embryo sac (dark blue) of Col-0 and *pTT1::BAR* ovules; bottom: 7-celled WT and *pTT1::BAR* embryo sacs. The WT embryo sac appears more elongated than the *pTT1::BAR* embryo sac. B. Plot depicting the embryo sac volume of WT and *pTT1::BAR* ovules at stages 3-I and 3-V of ovule development. Data points indicate individual ovules. Mean ± SD is shown. C. Mid-sagittal section of a stage 3-V *pTT1::BAR* ovule, also expressing the *FGR8.0* embryo sac marker and stained with the cell wall stain SR2200 (white). FGR8.0 embryo sac nuclei are labeled with *pEC1*.*1*::*NLS–3xDsRed2* (red) in the egg cell, *pLURE1*.*2*::*NLS–3xGFP* (yellow) in the synergid cells, and *pDD22::YFP* (yellow) in the central cell. Scale bars: 10 μm.

### Correct inner integument development is required for early embryo sac development and shape, but not for fertilization

In about 90 percent of *pTT1::BAR* ovules, embryo sac development was arrested around the mononuclear embryo sac stage while in about 10 percent of the cases, embryo sac development continued (Fig. 5B-C, Fig. 6A-B bottom panels, Table 2). The size of a 3-V *pTT1::BAR* embryo sac remained unchanged compared to the wild type (Fig. 5B), however, it appeared less elongated than in the wild type (Fig. 5A). The stage 3-V *pTT1::BAR* ovules appear to have regular embryo sac differentiation, i.e., the embryo sac has distinct synergids, an egg cell, a central cell, and antipodal cells (Fig. 5A). To confirm that embryo sac differentiation is comparable to wild type, we crossed the FGR 8.0 reporter line (Völz *et al*., 2013) with the *pTT1::BAR* line. The FGR 8.0 line carries several different reporters specific for the different cell types of the cellularized embryo sac (pLURE1.2::NLS-3xGFP, synergids; pEC1::NLS-3xDsRed2, oocyte; pDD22::YFP, central cell). In the ovules of the *pTT1::BAR FGR 8.0* line with a strongly reduced inner integument, we detected the expected reporter signals in the synergids, the egg cell and the central cell (Fig. 5C) indicating regular cell differentiation in the embryo sac. Taken together, our results suggest a role for the inner integument in the control of the early nuclear divisions of the female gametophyte and in the regulation of its shape.

**Fig. 6.**
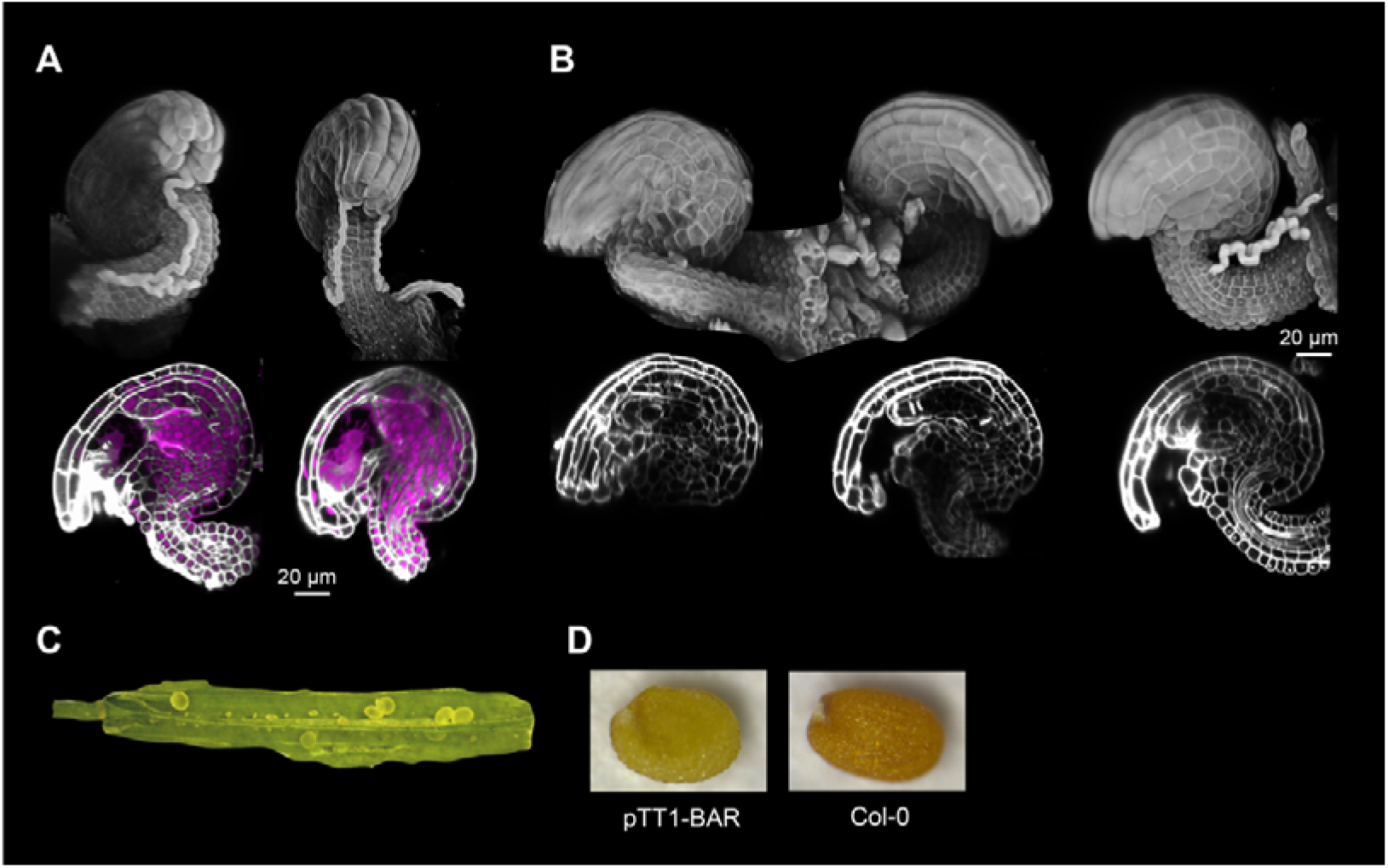
Pollen tube entry, pistil, and seed morphology of *pTT1::BAR* ovules. A. and B. 3D renderings of *pTT1::BAR* ovule confocal z-stacks (top) and their mid-sagittal sections (bottom) with SR2200-stained cell walls (white) and TO-PRO-3-stained nuclei (magenta). A. Few ovules that undergo pollen tube entry (top) possess stage 3-VI embryo sacs as seen from the embryo sac nuclei stained with TO-PRO-3 (bottom). B. Mature-looking *pTT1::BAR* ovules lacking pollen tube entry (top) due to embryo sac growth arrest at stage 2-V or 3-I (bottom). C. The *pTT1::BAR* pistil shows many degenerated ovules and very few seeds. D. Comparison of *pTT1::BAR* (left) and Col-0 (right) seed morphologies. Scale bars (A,B): 20 μm.

Pollen tube entry and fertilization depends on functional synergids and egg cells (Zhou and Dresselhaus, 2019; Sugi and Maruyama, 2023). Thus, we tested the functionality of *pTT1::BAR* ovules bearing mature embryo sacs. To this end, we examined pollen tube entry and fertilization in *pTT1::BAR* ovules (Fig. 6A-B). We observed approximately 2-6 ovules per pistil undergoing fertilization (2-10 percent). As expected, ovules associated with pollen tube entry contained differentiated embryo sacs (3-VI) (Fig. 6A). Very few ovules (0-2 per pistil) showed entry of two pollen tubes into the micropyle, suggesting an impairment of the polytubey block. Ovules from the same pistil lacking pollen tube attraction featured early arrested embryo sacs (Fig. 6B). The results suggest that about 2-10 percent of *pTT1::BAR* ovules, despite having a strongly reduced inner integument, undergo embryo sac differentiation and fertilization. Therefore, even though the inner integument is important in the control of nuclear divisions during the coenocytic phase of embryo sac development and its elongated shape, it is not essential for cellularization, cell differentiation, pollen tube entry and fertilization.

### Fertilized *pTT1::BAR* ovules undergo seed development

Next, we asked whether *pTT1::BAR* ovules could develop into seeds. Under our growth conditions, young developing siliques of Col-0 wild-type plants carried about 52 developing seeds (51.6 ± 3.5 seeds per silique (mean ± standard deviation, n=5 siliques from 5 different plants). We found that *pTT1::BAR* pistils exhibited a similar number of ovules but showed reduced seed set as they produced 3.8 ± 1.8 seeds per silique (mean ± standard deviation, n=3 plants, 20 siliques per plant). Thus, about 93 percent of the *pTT1::BAR* ovules remained unfertilized and degenerated, likely due to the early arrest of embryo sac development. About 7 percent of ovules underwent fertilization and developed into seeds (Fig. 6C). The shape of the *pTT1::BAR* seeds appeared dented and seed color was yellow rather than brown since they possess a highly reduced inner integument lacking the tannin-producing endothelium (ii1) (Fig. 6D).

The fertilized *pTT1::BAR* ovules were further assessed for endosperm and embryo development. Endosperm plays a crucial role during seed development. It has an important role in supplying nutrients and protecting the growing embryo (Doll and Ingram, 2022). During fertilization, one sperm nucleus fuses with the central cell, generating the primary endosperm cell which eventually develops into the endosperm. The *pDD22::YFP* central cell reporter also marks the non-cellularized endosperm (Steffen *et al*., 2007). We followed *pDD22::YFP* signal in fertilized *pTT1::BAR* ovules expressing the FGR 8.0 reporter. We observed strong *pDD22::YFP* expression in non-cellularized endosperm in *pTT1::BAR* ovules with a highly reduced inner integument (Fig. 7). We also observed expression of the *pPHE1::PHE1-GFP* early endosperm marker (Weinhofer *et al*., 2010) in fertilized *pTT1::BAR* ovules (Fig. S2). Our results suggest that the fertilized *pTT1::BAR* ovules undergo endosperm development even in the presence of a highly reduced inner integument.

**Fig. 7.**
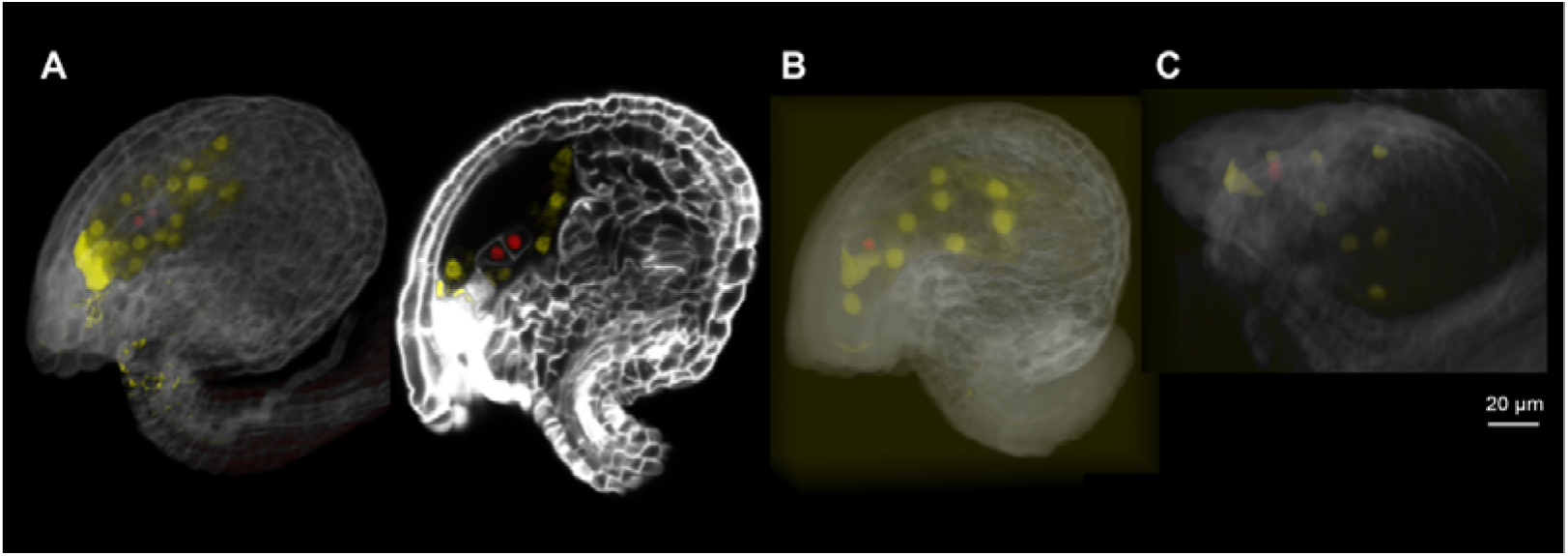
Fertilized *pTT1::BAR* ovules undergo normal seed development as demonstrated by the *FGR8.0* marker. (A-C) 3D renderings of confocal z-stacks and mid-sagittal section (A, right) of three *pTT1::BAR* ovule specimens expressing the *FGR8.0* marker and stained with the cell wall stain SR2200 (white). The 3D stacks have been adjusted for opacity, brightness, and contrast of the signals. The *pDD22::YFP* (yellow) marker labels the endosperm nuclei since they are derived from nuclear divisions of the central cell. The *pEC1*.*1*::*NLS–3xDsRed2* (red) labels the embryo nuclei. Scale bar: 20 μm.

We then examined embryo development in the developing seeds inside the *pTT1::BAR* siliques in comparison to that in wild-type (Fig. 8A-B). We observed that developing *pTT1::BAR* seeds carry developing embryos (Fig. 8B). We compared the cell volumes of 1-, 2-and 4-celled *pTT1::BAR* embryos to those of the correspondingly staged Col-0 wild-type embryos. The mean cell volumes of *pTT1::BAR* embryos at 1-, 2-, and 4-celled stages were 52%, 24.3%, and 24.5%, respectively smaller than those of the correspondingly staged wild-type embryos (Fig. 8C, Table 5). The mean embryo proper volume of globular stage *pTT1::BAR* embryos (2.69 × 10^4^ ± 1.31 × 10^4^ µm^3^ (mean ± SD); n = 6) was comparable and not statistically significantly different from those of wild-type embryos (2.33 × 10^4^ ± 0.88 × 10^4^ µm^3^ (mean ± SD); n = 33) (Fig. 8C). The results suggest that the reduction in the inner integument in *pTT1::BAR* ovules leads to an initial delay in cellular growth of embryos. However, by the globular stage the initial setback to growth is overcome and eventually the embryos sizes of the mutant and wild type are comparable.

**Fig. 8.**
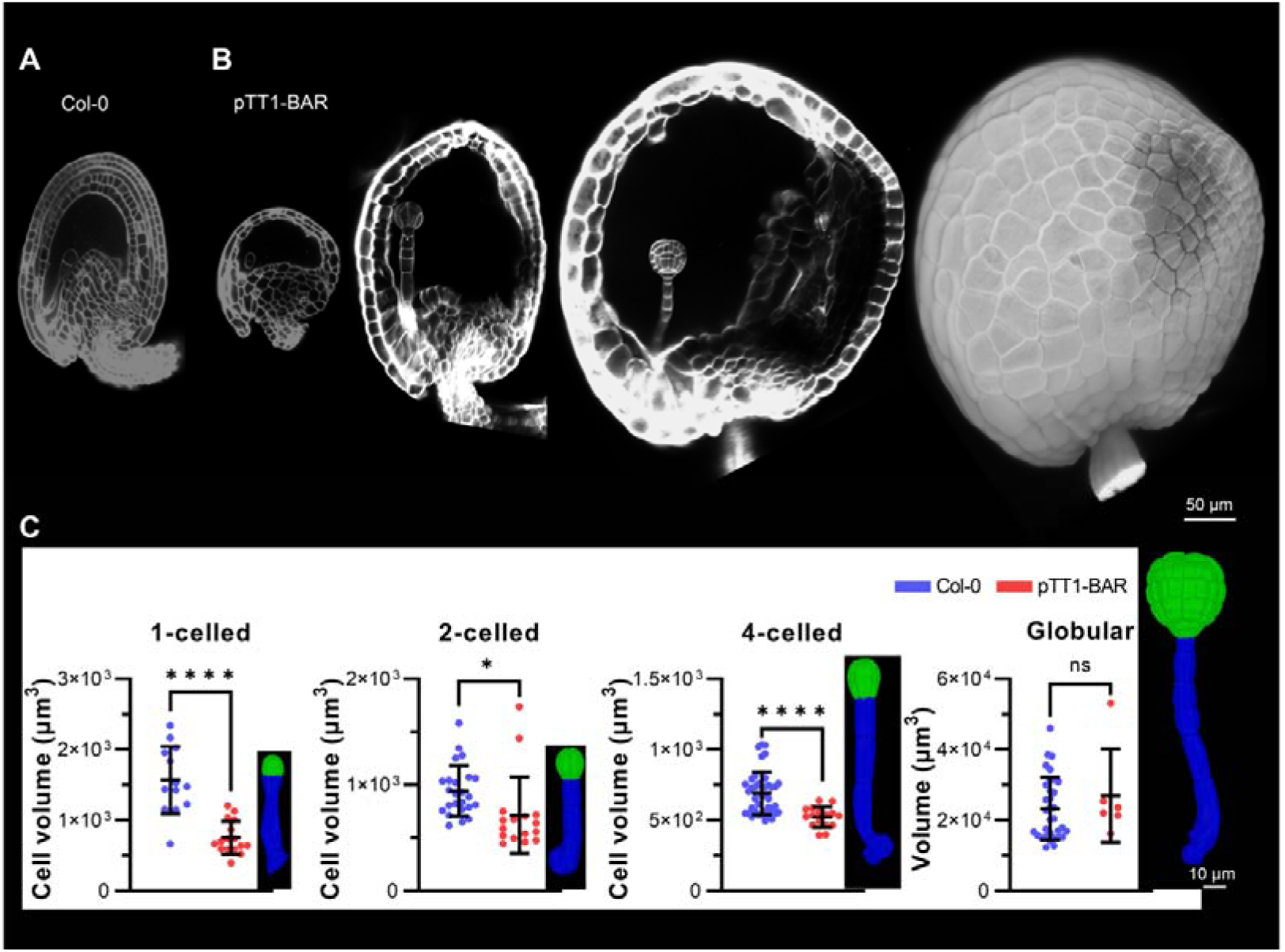
Fertilized *pTT1::BAR* ovules undergo normal embryo development. (A-B) Mid-sagittal sections of confocal z-stacks of ovules and seeds stained with the cell wall stain SR2200 (white). A. Col-0 WT ovule harboring a 1-celled embryo. B. *pTT1::BAR* ovule and seeds harboring (left to right) 1-celled, early globular, and late globular embryos. The rightmost image is a 3D rendering of the confocal z-stack of SR2200-stained *pTT1::BAR* seed. C. Plots depicting a comparison of cell volumes or volume between WT and *pTT1::BAR* embryos. The cell volume comparisons are performed for 1-celled, 2-celled, and 4-celled embryo stages, and embryo proper volume comparison is performed for globular stage embryos. The insets in black show the *pTT1::BAR* embryo phenotypes for the corresponding stages. Scale bar: 10 μm.

**Table 5.** Cell volumes of cells of 1-, 2-, and 4-celled *pTT1::BAR* and wild-type embryos.

|  | *pTT1::BAR* |  | Col-0 |  |
| --- | --- | --- | --- | --- |
| Embryo<br>proper stage | Cell volume (μm<sup>3</sup>)<br>(mean ± SD) | N* | Cell volume (μm<sup>3</sup>)<br>(mean ± SD) | N* |
| 1-celled | 752.3 ± 232.0 | 16 | 1568.0 ± 475.1 | 13 |
| 2-celled | 713.3 ± 358.6 | 8 | 942.9 ± 238.1 | 12 |
| 4-celled | 521.9 ± 72.7 | 4 | 691.1 ± 150.8 | 10 |

\*Number of embryo specimens scored.
Values for Col-0 were obtained from (Yoshida *et al.*, 2014)

Finally, we tested whether the seeds produced by the *pTT1::BAR* plants germinated and whether the respective seedlings could develop into mature plants. We found that except for a slight reduction in germination, seedlings developed into plants with apparently normal morphology aside from the expected defects in ovule and seed development (Fig. S3).

In summary, the results suggest that fertilized *pTT1::BAR* ovules undergo embryo and endosperm development, develop into viable seeds, which eventually give rise to mature plants despite a near-complete reduction in inner integument growth.

## Discussion

Here, we addressed the role of the inner integument in ovule and seed development by specifically interfering with the outgrowth of the inner integument. Our data indicate that expressing *BAR* in the innermost layer of the developing inner integument using a 3 kb *TT1* promoter fragment results in a near absence of the entire inner integument. In contrast, a previous study using a *BANYULS* (*BAN*) promoter fragment to drive *BAR* expression in the inner integument resulted in specific ablation of the postfertilization endothelium while leaving the other tissue layers of the inner integument in place (Debeaujon *et al*., 2003). What is the basis for the distinct effects on tissue elimination between the *pTT1::BAR* and *pBAN::BAR* lines? The later effects in *pBAN::BAR* lines are likely due to the mostly postfertilization expression of *BAN* (Debeaujon *et al*., 2003) while the *pTT1* promoter is already active during ovule development (Coen *et al*., 2019) (Fig. 1C). In terms of which tissues are eliminated, we postulate that the almost complete elimination of the entire inner integument in *pTT1::BAR* is related to BAR interfering with the mechanism that maintains the adaxial-abaxial polarity required for organ outgrowth and blade formation of lateral organs such as leaves and integuments (Waites and Hudson, 1995; McConnell and Barton, 1998; Villanueva *et al*., 1999; Eshed *et al*., 2001; Bowman *et al*., 2002; Kelley *et al*., 2009). In the case of *pBAN::BAR*, a fully developed inner integument is generated prior to the onset of *pBAN* expression, allowing targeted elimination of the endothelium during seed development. The absence of a completely eliminated inner integument in *pTT1::BAR* relates to the observation that the *pTT1* promoter fragment becomes stably active in all endothelial cells from stage 3-I onwards. The BAR toxin has been shown previously to cause cell growth arrest rather than cell death in trichome cells (Faden *et al*., 2019) and thus, similarly, the *pTT1::BAR* ovules show inner integument cell growth arrest but not cell death as demonstrated by the nuclear morphology. This is also supported by the observation that there are no drastic differences in the size of inner integument of *pTT1::BAR* ovules from stage 3-I to 3-VI (Fig. 2B, Table 1).

The majority of *pTT1::BAR* ovules exhibited an extremely reduced inner integument accompanied by an early block in embryo sac development. The molecular basis for this observation is currently unclear. The mechanism underlying the regulation of embryo sac development by the integuments is generally poorly understood and is likely to be complex (Qin *et al*., 2023). A role for auxin in ovule and seed development is well established (Shirley *et al*., 2019). For example, *TRYPTOPHAN AMINOTRANSFERASE OF ARABIDOPSIS* (*TAA*), a gene involved in tryptophan-dependent auxin production, is expressed in the early inner integument (Ceccato *et al*., 2013). In addition, the auxin efflux transporter PINFORMED 1 (PIN1) is present in the epidermis of the nucellus and inner integument, and PIN1 is required for early embryo sac development (Ceccato *et al*., 2013). In one model, auxin produced in the young inner integument is transported in a PIN1-dependent manner through the nucellar epidermis to the nucellar tip, where it is involved in early gametophyte development. Thus, a plausible explanation for the early embryo sac defects in the majority of the *pTT1::BAR* ovules could be a reduced auxin maximum at the nucellar tip due to a failure of regular auxin production in the severely reduced inner integument. Conversely, in the 10% of *pTT1::BAR* ovules that develop an embryo sac, auxin production in the inner integument may have been sufficient. It will be interesting to investigate the mechanism underlying the embryo sac defect in *pTT1::BAR* ovules in future studies.

Our data show that the 10% *pTT1::BAR* ovules with an embryo sac are fertile and develop into functional seeds despite the severely reduced inner integument. Interestingly, the functional embryo sac of a *pTT1::BAR* ovule generates the proper cell types and volume but its shape is altered compared to wild type. This finding suggests that the embryo sac obtains its elongate shape by mechanical influences exerted by the inner integument. The results further indicate that the inner integument is not required for fertilization. They are in line with findings that the synergids and the outer integument play essential roles in pollen tube guidance (Zhou and Dresselhaus, 2019; Mizuta *et al*., 2024).

Regarding seed development there is evidence that early embryo development depends on auxin supplied by the maternal integuments (Robert *et al*., 2018). Analysis of the expression patterns of the auxin sensor R2D2 (Liao *et al*., 2015) and the auxin response sensor *pDR5::GFP* (Friml *et al*., 2003) indicated an upregulation of auxin levels in maternal tissue surrounding the embryo and a localized upregulation of the auxin response at the embryo attachment site (Robert *et al*., 2018). Previous data indicated that the endothelium is not required for seed development (Debeaujon *et al*., 2003). Our data suggest that not only the endothelium but the entire inner integument is not essential for embryo and endosperm development.

Seed shape depends on endosperm turgor pressure and on differential growth patterns and mechanical forces in the two layers of the outer integument (Creff *et al*., 2023; Bauer *et al*., 2024). Our data suggest that the inner integument contributes substantially to seed shape, as it was markedly altered in *pTT1::BAR* seeds. Thus, the final seed shape depends on the interplay of several tissues, including the endosperm, inner and outer integuments.

In summary, our data provide evidence that the inner integument is required for embryo sac formation and seed shape, but is dispensable for embryo and endosperm development.

## Supplementary data

- Brief single-sentence description for each item

**Fig. S1.** Variability in morphology and inner integument reduction of stage 3-V *Arabidopsis thaliana pTT1::BAR* ovules.

**Fig. S2.** Fertilized *pTT1::BAR* ovules undergo normal endosperm development as demonstrated by the *PHE1::PHE1-GFP* marker.

**Fig. S3.** Seed germination and seedling development in Col-0 and *pTT1::BAR* line.

## Acknowledgements

We thank Rita Gross-Hardt for the FGR8.0 and Claudia Köhler for the pPHE1::PHE1-GFP seeds. We further thank Enrico Magnani for the 3kbProTT1:gTT1 plasmid. We acknowledge support by the Center for Advanced Light Microscopy (CALM) of the TUM School of Life Sciences.

## Authors’ contributions

TAM and KS designed the study. TAM performed the experiments. TAM and KS interpreted the results. KS secured funding. TAM and KS wrote the paper. All authors read and approved the final manuscript.

## Data availability

The Col-0 wild-type embryo 3D meshes were obtained from (Yoshida *et al*., 2014). The Arabidopsis wild-type 3D digital ovule dataset can be obtained from Biostudies (accession S-BSST475). The *pTT1::BAR* ovule dataset is available from Biostudies as well (accession S-BIAD1499).

