## Supplemental Figures for "The inner integument controls embryo sac development and seed shape in *Arabidopsis thaliana*": ModyandSchneitz_Supplement Kopie.docx

**Supplementary Figures S1-S3**

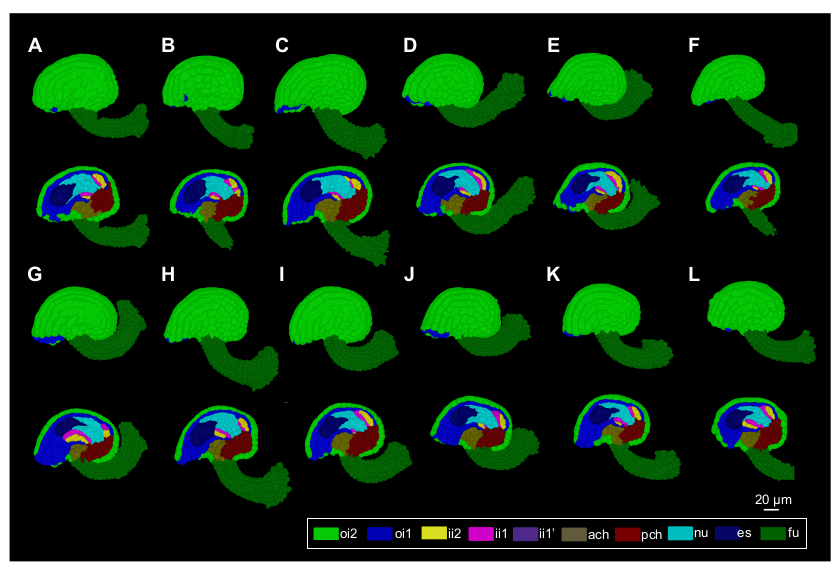

**Fig. S1.** Variability in morphology and inner integument reduction of stage 3-V *Arabidopsis thaliana pTT1::BAR* ovules. (A-L) A total of 12 stage 3-V *Arabidopsis thaliana pTT1::BAR* ovules with cell-type labels showing the tissue organization; top: 3D view and bottom: mid-sagittal section view. Abbreviations: ach, anterior chalaza; es, embryo sac; fu, funiculus; ii1, inner layer of inner integument; ii1’, parenchymatic layer of inner integument; ii2, outer layer of inner integument; nu, nucellus; oi1, inner layer of outer integument; oi2, outer layer of outer integument; pch, posterior chalaza. Scale bar: 20 μm.

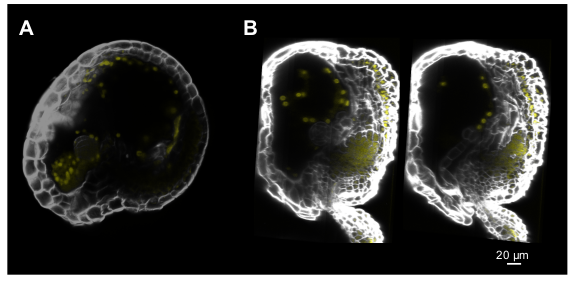

**Fig. S2.** Fertilized *pTT1::BAR* ovules undergo normal endosperm development as demonstrated by the *PHE1::PHE1-GFP* marker. (A-B) Mid-sagittal sections of confocal z-stacks of two *pTT1::BAR* ovule specimens expressing the *PHE1::PHE1-GFP* marker (yellow) and stained with the cell wall stain SR2200 (white). *PHE1::PHE1-GFP* marker (yellow) labels the endosperm nuclei. Scale bar: 20 μm.

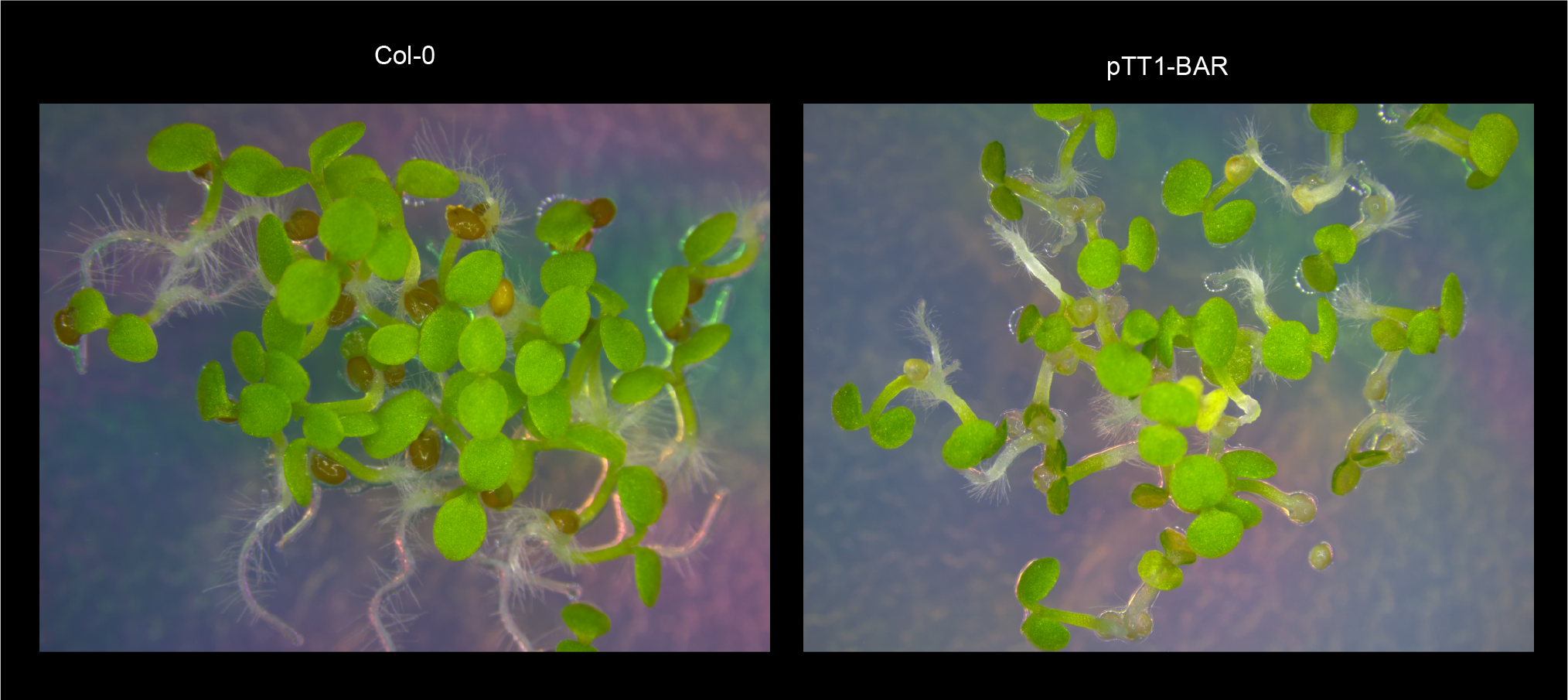

**Fig. S3.** Seed germination and seedling development in Col-0 and *pTT1::BAR* line. Images showing seedlings from (left) Col-0 wild-type and (right) *pTT1::BAR* line, 68 hours after seed stratification (has). Seeds produced by the *pTT1::BAR* plants germinated (with a slight reduction in germination) and respective seedlings could develop.
